# Exposure to hypomethylating agent 5-aza-2’-deoxycytidine (decitabine) causes rapid, severe DNA damage, telomere elongation and mitotic dysfunction in human WIL2-NS cells

**DOI:** 10.1101/834382

**Authors:** Caroline Bull, Graham Mayrhofer, Michael Fenech

## Abstract

**Background:** 5-aza-2’-deoxycytidine (5azadC, decitabine) is a DNA hypomethylating agent used in the treatment of myelodysplastic syndromes. Due to cytotoxic side effects dose optimization is essential. This study defines and quantifies the effects of 5azadC on chromosomal stability and telomere length, at clinically relevant dosages.

**Methods:** Human WIL2-NS cells were maintained in complete medium containing 0, 0.2 or 1.0μM 5azadC for four days, and analysed daily for telomere length (flow cytometry), chromosomal stability (cytokinesis-block micronucleus cytome (CBMN-cyt) assay), and global methylation (%5me-C).

**Results:** DNA methylation decreased significantly in 1.0 μM 5azadC, relative to control (p<0.0001). Exposure to 1.0μM 5azadC resulted in 170% increase in telomere length (p<0.0001), in parallel with rapid increase in biomarkers of DNA damage; (micronuclei (MN, 6-fold increase), nucleoplasmic bridges (NPB, a 12-fold increase), and nuclear buds (NBud, a 13-fold increase) (all p<0.0001). Fused nuclei (FUS), indicative of mitotic dysfunction, showed a 5- and 13-fold increase in the 0.2μM and 1.0μM conditions, respectively (p = 0.001) after 4 days.

**Conclusions:** These data show that (i) clinically relevant concentrations of 5azadC are highly genotoxic; (ii) hypomethylation was associated with increased TL and DNA damage; and (iii) longer TL was associated with chromosomal instability. These findings suggest that lower doses of 5azdC may be effective as a hypomethylating agent, while potentially reducing DNA damage and risk for secondary disease.

## Introduction

DNA methylation is essential for gene transcription control, possibly evolving from the need to silence genes of parasitic or viral origin (1). Dysregulation of the epigenome is associated with neoplastic changes and tumorigenesis (1, 2). Hypermethylation contributes to the aetiology, and a worsening of clinical symptoms in myelodysplastic syndromes (MDS) and acute myeloid leukemia (AML). A bone marrow stem cell transplant offers the greatest chance of cure, but this is not a viable option for many patients, particularly the elderly. In some cases hypomethylating drugs such as 5-azacytidine and decitabine (5-aza-2’-deoxycytidine (5azadC)), used alone or in combination with other chemotherapeutics, can extend survival time and improve quality of life (3). These regimens are, however, highly cytotoxic and can lead to severe (acute) side effects. DNA damage resulting from treatment can also cause longer term effects such as impaired immune function and development of secondary cancers (3, 4). It is essential to optimize treatment protocols to maximise efficacy, while minimizing genotoxic side effects, and secondary health impact.

5azadC is an analogue of the pyrimidine nucleoside cytidine, in which the carbon at position 5 is replaced with nitrogen. It has been used therapeutically for the treatment of all MDS subtypes since FDA approval in 2004 (3–5). 5azadC binds irreversibly (covalently) with DNA methyltransferase 1 (DNMT1), resulting in (i) adduct formation within the DNA sequence which potentially obstructs DNA synthesis, and (ii) inhibition of DNMT1 from catalysing further methylation reactions. The hypomethylating effect becomes more pronounced after several cell divisions, as numbers of free DNMT1 molecules are gradually depleted (6). The net effect is to reduce/reverse the degree of DNA methylation, thus reactivating aberrantly silenced genes, such as tumor suppressors (6). A review of decitabine dosage protocols for MDS patients showed a range from 15-500 mg/m^2^ infused over 1-6 hours, with some as long as 120 hours. Resulting plasma concentrations ranged from 0.12 – 5.6 μM (4). In a cohort of patients taking oral decitabine (in combination with tetrahydrouridine (THU)) a >75% reduction in DNMT1 in peripheral blood mononuclear cells (PBMC) was recorded, in parallel with (LINE-1) CpG methylation reduction of approximately 10% (5).

Previous studies to determine the degree of cytotoxicity and DNA damage caused by 5azadC have measured the extent to which H2AX, a key ‘first responder’ in a DNA breakage damage response, is phosphorylated to generate γH2AX (7). While informative, this method is specific to DNA breaks and does not capture the different forms of chromosomal instability induced by hypomethylation and DNA replication stress. The cytokinesis-block micronucleus cytome (CBMN-cyt) assay is a comprehensive, robust diagnostic tool for detecting and quantifying several chromosomal instability events such as micronuclei, nucleoplasmic bridges and nuclear buds. The micronucleus (MN), has been internationally validated as a risk marker for cancer risk and cardiovascular disease mortality (8). To our knowledge the impact of 5azadC has not previously been studied using the CBMN cytome assay. However a previous study showed that 5azadC can induce MN due to malsegregation and loss of chromosomes 1, 9, 15, 16 and Y (9). Furthermore, an additional measure was examined in the present study, examining frequencies of the novel biomarker FUSED nuclei (FUS). FUS are indicative of failed chromatid separation, or telomere end fusions, and are known to be sensitive to changes in methylation status (10).

An additional biomarker for examining genome stability, and disease risk, is telomere length (TL). Telomeres are complex nucleoprotein structures that cap chromosome ends, protecting the coding gene region from degradation during cell division (11). Comprised of a hexamer repeat sequence which lacks CpGs (TTAGGGn), telomeric DNA has no substrate for DNMT enzymes, and as such are unmethylated. The impact of changes to methylation status on telomeres varies, with conflicting results reported depending on cell type. Several studies have shown that TL increases under hypomethylating conditions (12–14), with evidence suggesting this effect is mediated through changes at the subtelomere, a heavily methylated region located between the telomere and coding DNA (14). As increased risk for many cancers has now been associated with longer TL (15–20), it is not appropriate to conclude that telomere elongation is intrinsically healthier, or provides greater stability. Ideally TL measures should be conducted in parallel with measures of chromosomal stability to distinguish between telomere lengthening that promotes chromosomal stability from that which does not. Our own previous findings have shown that longer telomeres induced under hypomethylating conditions are dysfunctional, and associated with increased chromosomal instability and DNA damage (10, 13).

The aim of the current study was to examine the hypothesis that exposure to 5azadC, within a clinically relevant range, would cause an increase in both chromosomal instability, and telomere length (TL). A key aim was to define and quantify the types of DNA damage induced. To test this, human WIL2-NS cells were cultured in the presence of 0, 0.2 or 1.0 μM 5azadC for 4 days. Samples were analysed daily for growth (nuclear division index), viability, telomere length, DNA methylation status, and a panel of biomarkers of DNA damage using the CBMN-cyt assay.

## Materials and Methods

### Study Design

Human WIL2-NS (B lymphoblastoid) cells (American Type Culture Collection (ATCC); CRL-8155) were maintained in culture for 4 days, in complete medium containing 0, 0.2 or 1.0 μM 5azadC. Cells were sampled daily and assessed for growth and viability, telomere length (TL) by flow cytometry, global methylation status (ELISA), chromosomal instability and DNA damage (cytokinesis block micronucleus cytome (CBMN-Cyt) assay). WIL2-NS was selected as an ideal (and proven) model to examine DNA damage events. The p53-deficient status of the cells allows for observation of substantial genome damage events without excessive cell death through apoptosis.

### Cell culture

Medium was prepared using RPMI 1640 (Sigma R0883), supplemented with 5% foetal bovine serum (FBS) (Thermo, Australia), 1% penicillin/streptomycin (Sigma P4458) and 1% L-Glutamine (Sigma G7513). 5azadC (5 mg in powder form, Sigma A3656) was dissolved in 1 mL 1x phosphate buffered saline (PBS) to form a 5 mg/mL stock solution with a final concentration of 21.9 mM, sterilised by filtration, and stored in aliquots at −20°C. Pilot studies were conducted with cells cultured in CM containing 5azadC within a clinically relevant range; 0.2, 1, 5 and 10 μM. Results showed 0.2 and 1.0 μM were optimal concentrations for maintaining a growth/survival rate sufficient to perform experimental assays up to four days. This timeframe was selected based on previous studies which had shown this to be sufficient to produce large reductions in global DNA methylation, as well as the development of micronuclei (21–23). WIL2-NS cells were thawed from liquid nitrogen storage, washed twice in PBS and grown in complete medium (CM) for seven days at 37°C in a humidified atmosphere with 5% CO_2_. Cells were split into the three different 5azadC conditions (day 0 time point), and cultured for a further 4 days. Cultures were maintained at 90 mL volume in a 75 cm^2^ vented-cap culture flask (Becton Dickinson, Australia). Duplicate flasks were established at day 0 to cater specifically for each sample day (days 1, 2, 3 and 4), totalling eight flasks per condition. Initial seeding concentration of each pair of flasks was adjusted, based on pilot growth data, to ensure the required numbers of viable cells at each sample point, while minimising overcrowding, nutrient depletion and pH imbalance. Each day, two flasks were removed for analysis, leaving the remaining flasks undisturbed until their respective sample day.

### Measurement of telomere length (TL)

Telomere length (TL) was measured in cells at G1, using the flow cytometric method described previously (24). In brief, fixed and permeabilised cells were labelled with an 18mer FITC-conjugated peptide nucleic acid (PNA) probe complementary to the telomere repeat sequence, using kit K5327 (Dako, Denmark), and counterstained with propidium iodide (PI) to measure DNA content and identify cell cycle stage. As a reference, cells from a tetraploid line with long telomeres (cell line 1301; accession number 01051619, European Collection of Cell Cultures, UK) were included in all tubes and used to calculate the relative TL in lymphocytes in the test samples. Each sample was prepared in paired tubes. For the purpose of quantifying background fluorescence of both the sample and reference cells one tube was incubated in hybridisation mixture with the PNA probe, while the paired tube was incubated in hybridisation mixture only. TL and DNA content measurements were acquired using a FACSCalibur flow cytometer (Becton Dickinson) and analysed using BD CellQuestTM Pro software (v5.2). A mean FITC fluorescence value was obtained for the sample and 1301 cells in the labelled and unlabelled samples by gating specifically at G0-1 phase of the cell cycle. TL of sample cells relative to that of 1301 cells was then calculated, with correction for ploidy (DNA index) of the different cell populations. Baseline TL was mean of n=20 (CV 8%), all points thereafter were a mean of n=10 replicates per treatment per time point.

### Determination of chromosomal instability and damage using CBMN-Cyt assay

Chromosomal damage was assessed using a modified form of the CBMN-Cyt assay, the standard protocol for which has been described in detail elsewhere (8). In brief, duplicate 500 μL subcultures were established from cells harvested each day, at a concentration of 0.3 x 10^6^ viable cells / mL. Cytochalasin B (CytB) (4.5 μg / mL; Sigma, Australia) was added for the purpose of blocking cytokinesis, and cells were harvested after 24h by cytocentrifugation (Shandon Scientific, Cheshire, England). Slides were air-dried for 10min, fixed and stained using the commercial kit ‘Diff Quik’ (Lab Aids, Narrabeen, Australia). Slides were scored by one person (CB) using established criteria described by Fenech (8). In all, 1000 binucleated (BN) cells per duplicate culture (total 2000 BN cells per treatment per sample point) were scored for frequency of validated biomarkers of chromosomal instability (CIN) and damage, ie. BN cells containing micronuclei (MN), nucleoplasmic bridges (NPB) or nuclear buds (NBud). In addition, cells displaying ‘fused’ (FUS) nuclear morphologies, were scored in this study using criteria previously defined (10). Cytotoxic and cytostatic effects were determined in 500 cells per duplicate culture by scoring the rate of necrosis, apoptosis and nuclear division index (NDI). NDI was calculated as NDI = (M1 + 2M2 + 3M≥3)/*N*, where M1, M2 and M≥3 represent the number of cells with 1, 2 or ≥3 nuclei, and *N* is the total number of viable cells scored (i.e. excluding necrotic and apoptotic cells).

### DNA isolation and global DNA methylation

DNA was isolated using a DNEasy blood and tissue kit (Qiagen, Cat no. 69506) as per manufacturer’s instructions. Methyl-cytosine content, as a percentage of total cytosine content, was estimated using the MethylFlash Methylated DNA Quantification Kit (Colorimetric) (Epigentek, USA, Catalog No P-1034), following manufacturer’s instructions. The percentage of methylated cytosines in each sample was estimated using the following formula:

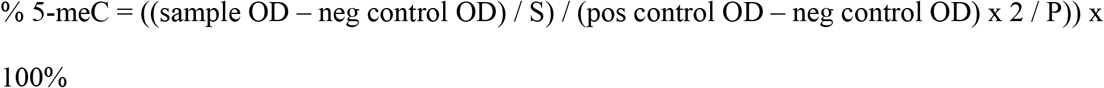

where OD is the optical density reading for each well (at 450 nm), ‘neg’ is negative control, S is the amount of input sample DNA in ng (ie. 100 ng was added per well), ‘pos’ is positive control, and P is the amount of positive control in ng. The numeral 2 in the denominator is required to normalise 5-meC in the positive control to 100%, as the sample provided in the kit is only 50% methylated. Samples were analysed in triplicate.

### Statistical analyses

Data are shown as mean ± SEM for all figures and tables. Two-way analysis of variance (ANOVA) was used to compare the effects for treatment ([5azadC]), time, and the interactive effect of these two factors. Pair-wise comparison of significance was determined using Bonferroni post-hoc test. Significance was accepted at p < 0.05. The area under the curve (AUC) for the relationship between changes in TL with time, was measured to obtain a total effect measure. Area under the curve (AUC) data represents the net area of the region in the xy plane bounded by the graph, where x is time (days) and y represents the parameter in question. All statistical analyses were performed using Graphpad PRISM 4.0 (GraphPad Inc., San Diego, CA).

## Results

### 5azadC exposure causes reduced nuclear division index and increased necrosis

The percentage of viable cells reduced in the 1.0 μM condition from 98% at day 0 to 40.6% at day 4 (p trend < 0.0001) (Fig 1A). Cells with necrotic morphology increased in both the 0.2 and 1.0 μM 5azadC conditions, to approximately 50% cell death in the 1.0 μM cultures by day 4 (p trend = 0.003) (Fig 1B). Consistent with these observations, nuclear division index (NDI) reduced significantly in both 5azadC concentrations (Fig 1C).

**Fig 1.**
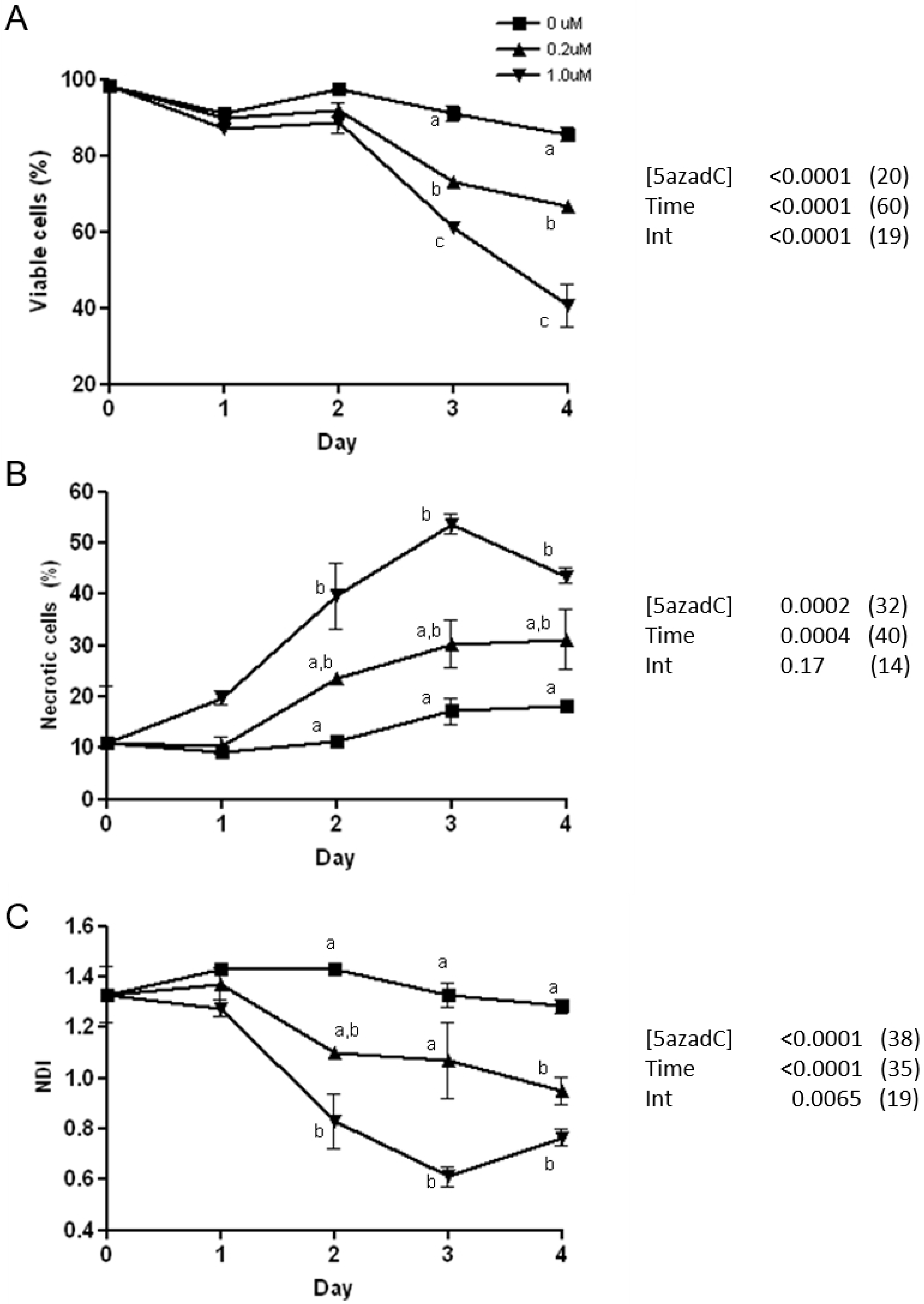
Impact of 5azadC on human WIL2-NS cells cultured in complete medium containing 0, 0.2 or 1.0 μM 5azadC over 4 days; (A) cell viability (%), (B) necrosis (%), and (C) nuclear division index (NDI). (N = 2. Error bars indicate SEM. Groups not sharing the same letter at each time point differ significantly from each other, as measured by the Bonferroni post-hoc test. Data tables alongside each graph indicate ANOVA p values for [5azadC], time and their interaction; figures in parentheses represent the degree of variance explained by each factor (%)).

### 5azadC causes telomere lengthening and global hypomethylation

Exposure to 5azadC caused significant, dose dependent, increase in TL (Fig 2A). In the 1.0 μM cultures TL increased by 172% from 19.2 ± 0.4 (mean ± SEM) at day 0, to 33.1 ± 0.6 at day 1, 156% longer than TL of cells maintained in 0 μM 5azadC for the same period (21.2 ± 0.5). At days 3 and 4, TL in cells cultured in 1.0 and 0.2 μM 5azadC were ~150% and ~130% greater, respectively, than that of cells cultured over the same time period without 5azadC (Fig 2A). [5azadC] accounted for 25.3% of variance in TL (p < 0.0001), 32.7% was due to time (p < 0.0001), with 30% attributable to the interaction of both factors (p < 0.0001). Area under the curve for TL versus time of cells cultured in 1.0 μM was 111, 28% greater than that of untreated cells (AUC 87), and 5% greater than TL of cells grown in 0.2 μM 5azadC (AUC 91).

**Fig 2.**
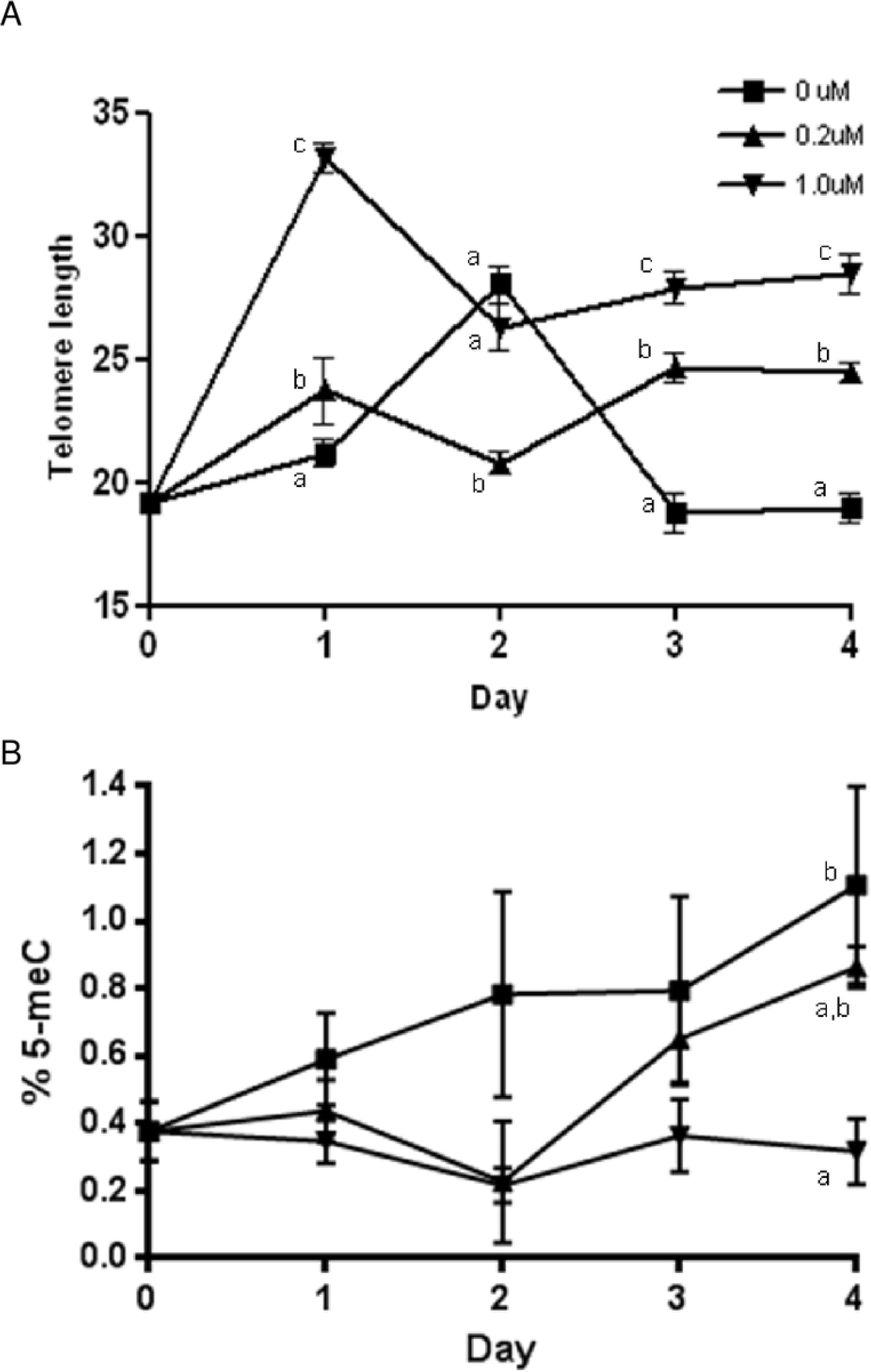
WIL2-NS cells grown in complete medium containing 0, 0.2 or 1.0 μM 5azadC over 4 days. (A) Telomere length (expressed relative to that of reference cell line, 1301 (%), n = 20 at day 0, n = 10 for all time points thereafter). (B) Global methylation status (% 5-meC, n = 3). (Error bars indicate SEM. Points not sharing the same letter at each time point differ significantly, as measured by the Bonferroni post hoc test).

In untreated (control) cells, and those exposed to the 0.2 μM 5azadC, global DNA methylation increased progressively with time, possibly due to supraphysiological concentrations of folic acid and methionine in RPMI. In the 1.0 μM 5azadC cultures methylation decreased resulting in significant differences in methylation status between conditions, in a dose-and time-dependent manner. In fact, 24% of variance in %5-meC was attributable to [5azadC] (p = 0.002), 18% was due to time (p = 0.04), with 25.3% of variance in methylation being explained by the interaction of [5azadC] with time (p = 0.3, not significant) (Fig 2B).

### 5azadC causes a rapid increase in DNA damage

Data generated with the CBMN-cyt assay showed exposure to 5azadC caused a time and dose-dependent increase in the frequency of biomarkers of chromosomal instability and DNA damage; micronuclei (MN), nucleoplasmic bridges (NPB), nuclear buds (NBuds), and fused nuclei (FUS) (8, 10). DNA damage is calculated based on the frequency of binucleated cells (BN), per 1000 BN, which contain one or more of each damage biomarker.

MN represent biomarkers of chromosome breakage or loss (8). The frequency of MN (per 1000 BN) in the 1.0 μM 5azadC cultures increased 6-fold over the course of the 4-day study; from 17 ± 1.0 at day 0, to 103 ± 14 at day 4. Over the same time period MN in cells in 0.2 μM 5azadC increased 4.5-fold to 79.5 ± 0.5 at day 4, while the frequency of MN in control cultures (0 μM 5azadC) reduced to 6.5 ± 1.5 at day 4 (Fig 3A).

**Fig 3.**
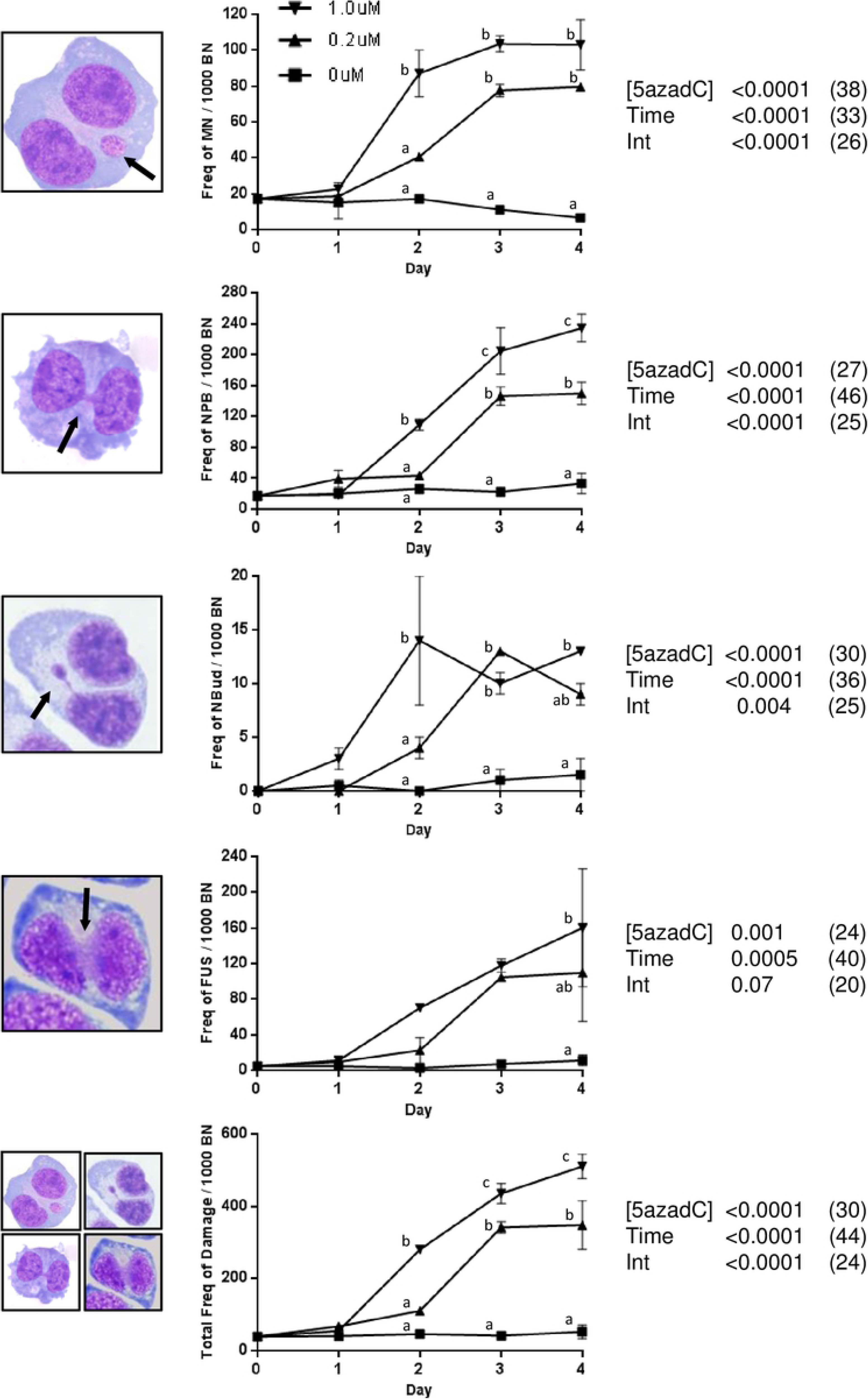
DNA damage in WIL2-NS cells grown in complete medium containing 0, 0.2 or 1.0 μM 5azadC for 4 days. Graphs represent the frequency of binucleated (BN) cells displaying one or more DNA damage biomarker per 1000 BN. (A) Micronuclei (BN-MN); (B) nucleoplasmic bridges (BN-NPB); (C) nuclear buds (BN-NBud); (D) ‘fused nuclei’ (BN-FUS) morphologies; and (E) total damage (frequency of BN cells with one or more MN and/or NPB and/or NBud and/or FUS per 1000 BN). (N = 2 cultures, 1000 BN scored per culture. Error bars indicate SEM. Groups not sharing the same letter at each time point differ significantly from each other, as measured by the Bonferroni post-hoc test. Data tables indicate ANOVA p values for [5azadC], time and their interaction; figures in parentheses represent the degree of variance explained by each factor (%)).

Similar effects were observed for BN containing NPB, with day 4 frequencies increased 12-fold (from 17 ± 3 at day 0 to 234 ± 18), 7.8-fold (to 149.5 ± 15), and 2-fold (to 33 ± 13) for the 1.0, 0.2 and 0 μM cultures, respectively (Fig 3B). NPB result from dicentric chromosomes caused by mis-repair of double strand DNA breaks, or telomere end fusions arising from telomere shortening and/or telomere dysfunction. NPB can indicate loss of telomeres due to DSB. These detached telomeric acentric chromosome fragments may still be detected with PCR or Southern blot TL assays. Only functional or visual assays, however, such as CBMN or FISH, can demonstrate that while telomeric DNA may be present, it is not located at chromosome ends, and thus is no longer protective or functional.

NBuds represent gene amplification, often arising due to breakage-fusion-bridge cycling. The frequency per 1000 BN cells containing one or more NBud increased from zero at day 0, to 13 ± 0 at day 4 in the 1.0 μM culture, and 9 ± 1 in the 0.2 μM condition (Fig 3C).

A 13-fold increase in BN cells with FUS nuclear morphologies was observed in the 1.0 μM condition, from a baseline of 38.5 ± 6.5 per 1000 BN, to 510 ± 34 at day 4. In the 0.2 μM condition FUS cells increased 5-fold to 347 ± 68, while frequencies in the control condition remained stable (Fig 3D). Examples of the FUS morphologies observed with 5azadC exposure are provided in Fig 4.

**Fig 4.**
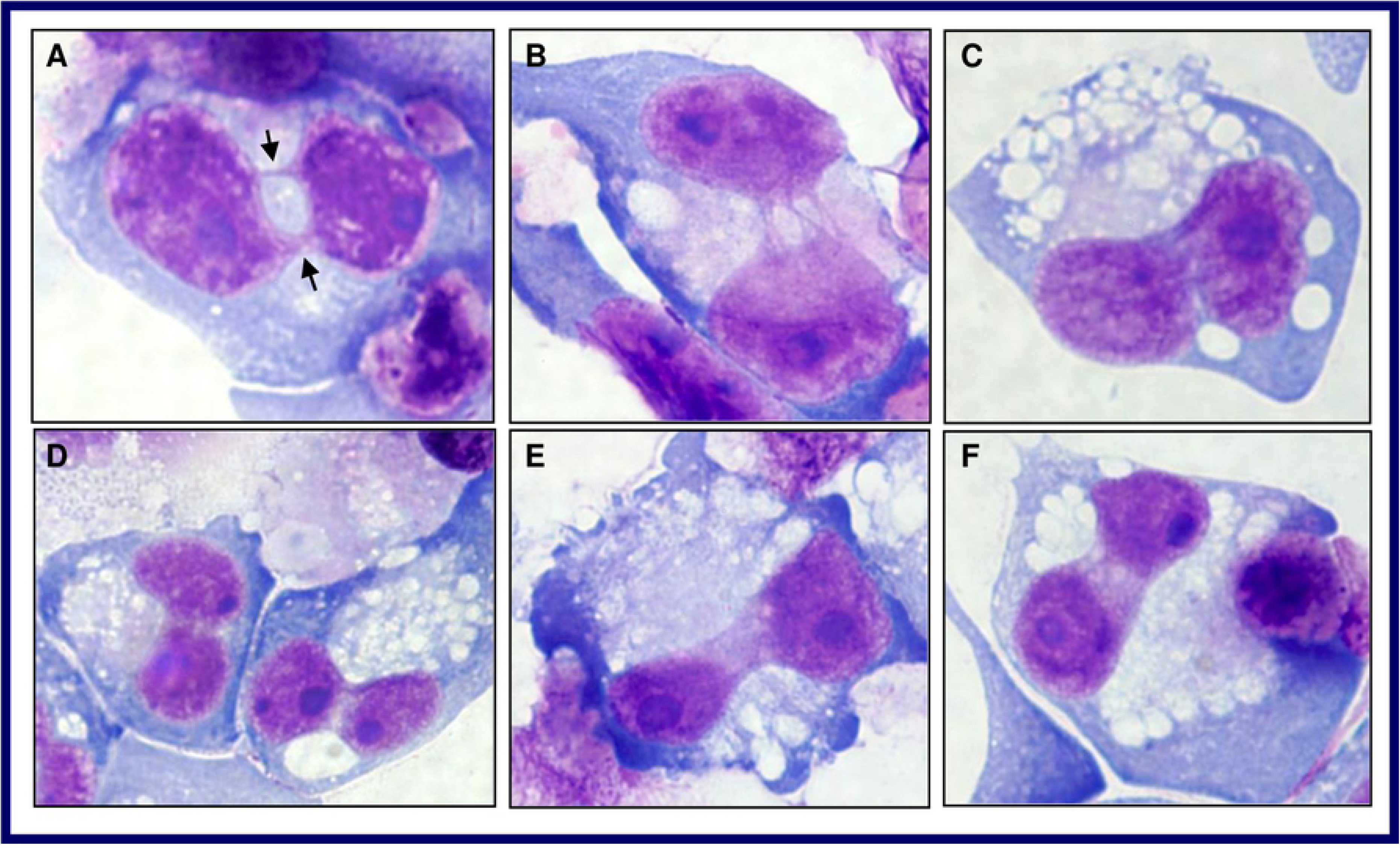
Representative photomicrographs of nucleoplasmic bridges (NPB) and fused (FUS) morphologies in WIL2-NS cells cultured for four days in medium containing 1.0 μM 5azadC, blocked at the binucleate (BN) stage using cytochalasin-B. (A) An example of a BN cell displaying two clearly discernible NPBs; (B) an example of a BN cell displaying four discrete NPBs; (C-F) Examples of BN cells with FUS morphologies (1000x magnification).

When examining the total combined frequency of BN cells containing a damage biomarker (ie. one or more MN and/or NPB and/or NBud and/or FUS morphology)), 30% of variance was attributable to [5azadC] (p < 0.0001), 44% was attributable to time (p < 0.0001), and 24% of the observed variance could be explained by the interaction of both factors (p < 0.0001), leaving only 2% of the variance unexplained (Fig 3E).

## Discussion

5azadC (decitabine), is used therapeutically to treat myelodysplastic conditions with dosage protocols resulting in plasma concentrations in the range of 0.12 – 5.6 μM (4). The hypothesis for the current work, that exposure to 5azadC, within a clinically relevant range, would cause an increase in both chromosomal instability, and telomere length (TL), was supported by the data.

Strong dose-dependent increases were observed for the frequency of BN cells containing one or more biomarker of DNA damage (MN, NPB, NBud, or FUS). The increase in MN indicates chromosome loss or dysfunction and/or double strand DNA breaks (DSB) (8). These observations are consistent with previous work on the effects of 5azadC showing that only a few hours exposure caused single and double strand breaks, together with decondensation of genetically inactive chromatin (25). 5azadC is a robust inducer of γ-H2AX, an early marker of DSB, activation of DNA repair proteins, and DNA fragmentation (7, 26, 27). Interestingly, the reduction of DNMT1 caused by 5azadC also impairs the cell’s capacity to respond to damage, as DNMT1 is required to co-localise with γ-H2AX at sites of damage. 5azadC also blunts the p53 and CHK1 responses in HeLa and HCT116 cells, further impacting an effective repair response (26). The concentration at which 5azadC-induced γ-H2AX foci and DNA strand breakage in these studies was at or below 1 μM (26), consistent with those used in the present study. These reflect clinically relevant plasma concentrations which range between 0.12 and 5.6 μM (4). Furthermore it was shown 5-azacytidine treated lymphocytes have been shown to exhibit distinct under-condensation in the heterochromatic regions of chromosomes 1, 9, 15, 16 and Y, and increases in the frequency of their loss via MN (9). These features are similar to the ICF syndrome in which DNMT 3B is defective (28).

NPBs are formed by DNA mis-repair following a DSB, or chromosome end fusion when telomeres become shortened and/or dysfunctional (8, 28–30). These fusions result in the formation of dicentric chromosomes which then present as NPBs in binucleated (BN) cells, suspended at telophase with the chromatids unable to separate. In cells that have not been chemically blocked at the BN stage, dicentric chromosomes will eventually break unevenly at anaphase, resulting in each of the daughter nuclei receiving abnormal gene dosage. The resulting uncapped ends are likely to fuse again with each other, leading to increasing levels of genomic disarray and instability. This is the breakage-fusion-bridge (BFB) cycle, an early event after telomere loss which leads to amplification of genes (including oncogenes), and altered gene dosage in daughter cells (8, 31). The increase in NBuds observed in cultures containing 5azadC is indicative of amplified genes or unresolved DNA repair complexes being actively ejected from the cell. Previous work has shown that, in 5azadC treated cells, BFB cycles can last many generations after the initial sister chromatid fusion (25, 31). The present findings are also consistent with those of Gisselsson *et al* (2005), who found an increased frequency of NPB in ICF cells that lack genes encoding DNMT3b (28).

In addition to the standard CBMN-cyt assay biomarkers, the frequency of novel FUSED (FUS) morphologies was also scored (8, 10). Previous findings showed that FUS nuclei increased significantly in WIL2-NS cells and primary lymphocytes cultured under folic acid-deficient (hypomethylating) conditions (10). Dual-colour fluorescence *in situ* hybridisation (FISH) analysis revealed centromeric DNA was present in the fusion structures between the nuclei. A mechanistic model was then proposed whereby hypomethylation alters the topology of the binding sites of key enzymes required for nuclear division (including Topoisomerase II and CENP-B), resulting in mitotic disruption (10, 32). As predicted, frequencies of FUS in the 5azadC treated cells increased significantly (13-fold) over 4 days, lending further weight to use of this biomarker as an indicator of hypomethylation-induced mitotic dysregulation.

Telomere elongation also occurred in a dose-dependent manner within 24 hours of 5azadC exposure, and was maintained to the completion of the study at day 4. There are conflicting findings with respect to TL with 5azadC treatment. In the breast cancer cell line 21NT TL increased (33), whereas in chronic leukemia cell lines telomeres shortened (34). The present findings are consistent with those of previous work in WIL2-NS cultured under (methyl donor) folic acid-deficient conditions, where the longest TL was recorded in the most severely FA-depleted condition (13). This observation is also consistent with the work of Gonzalo et al (2006), in which mouse embryonic stem cells genetically deficient in DNMT were found to have significantly longer telomeres than wild-type (12, 14). Epigenetic changes also affected TL in a panel of neoplastic cells, with significant negative associations between TL and methylation status at pericentromeric and subtelomeric sites (35). These authors demonstrated that subtelomeric hypomethylation was strongly associated with increased TL, and that this effect was independent of the expression, or activity, of the telomerase enzyme (35).

## Conclusions

The impact of 5azadC has not previously been examined using the comprehensive panel of DNA damage biomarkers included in the CBMN-Cytome assay. It is also essential to consider telomere length data in the context of chromosomal stability; as such, the findings presented here are novel. While consistent with previous observations of cytotoxicity, until such time as these assays can be replicated in other models the findings reported here are relevant only to the WIL2-NS cell line. We can conclude that, in this model, 5azadC at clinically relevant dosages induces hypomethylation and increased telomere length, BFB cycling, altered gene dosage and nuclear budding, chromosome breakage, mitotic dysregulation and high levels of DNA damage. These data also suggest that lower doses of 5azadC may provide a clinically efficacious level of hypomethylation, while minimising cytotoxic and genotoxic side effects, and risk for future secondary cancers caused by induced chromosomal instability.

## Author contributions

CB carried out all experimental work (cell culture, TL, CBMN-cyt assay, DNA methylation), data analysis and drafted the manuscript. GM, MF and CB contributed to the initial study concept and design, and interpretation of data. All authors had full access to the data. MF and GM both provided critical revision of the text and figures. Sadly, GM passed away in October 2016, thus CB and MF read and approved the final manuscript.

## Acknowledgements

The authors gratefully acknowledge the technical support of Ms Maryam Hor.

